# Inhibition of CKAMP44 attenuated seizure activity via protein phosphatase 3 regulatory subunit B-mediated GluA1 phosphorylation and synaptic transmission

**DOI:** 10.64898/2026.04.21.719815

**Authors:** Lihong Huang, Shiyu Chen, Haokun Guo, Hui Zhang, Liang Wang, Xiaoqin Wang, Yiming Guo, Shiyun Yuan, Jing Luo, Yang Lü, Weihua Yu

**Author notes:** Correspondence: Prof. Weihua Yu, Institutes of Neuroscience, Chongqing Medical University, Chongqing 400016, China Add: No.1 Yixueyuan Road, Yuzhong District, Chongqing 400016, China; Prof. Yang Lü, Department of Geriatrics, The First Affiliated Hospital of Chongqing Medical University, Chongqing 400016, China Add: No.1 Youyi Road, Yuzhong District, Chongqing 400016, China. These authors are co-corresponding authors. Authors contributed equally and are co-first authors of the study (Lihong Huang, Shiyu Chen, Haokun Guo).

## Abstract

Temporal lobe epilepsy (TLE) is a complex neurological disorder characterized by spontaneous recurrent seizures and its underlying mechanism remains elusive. This study aimed to investigate the role of cystine-knot AMPAR modulating protein 44 (CKAMP44) in the pathological process of TLE and its potential as a therapeutic target using kainic acid (KA)-induced epilepsy mouse model of TLE. Our results showed that CKAMP44 protein and mRNA expression was significantly increased and primarily localized to neurons during the chronic phase of TLE. Nkx2-1 regulated the transcription of CKAMP44 in the hippocampus brain tissues of KA-induced TLE mice. Inhibition of CKAMP44 suppressed seizure susceptibility and severity in the KA-induced epilepsy mice via behavioral and local field potential monitoring. Furthermore, inhibition of CKAMP44 decreased frequency and amplitudes of spontaneous excitatory postsynaptic currents indicating that the excitatory synaptic transmission was reduced in an in vitro epilepsy model. Mechanistically, inhibition of CKAMP44 specifically upregulated the membrane surface expression of GluA1 and the phosphorylation level of GluA1-ser831 by downregulating protein phosphatase 3 regulatory subunit B(PPP3r2) expression. Overexpression of PPP3r2 downregulated the phosphorylation level and surface expression of GluA1, which ultimately exacerbated the seizure activity suppressed by CKAMP44 knockdown. Collectively, our results indicate that CKAMP44 may be a potential therapeutic target for the treatment of TLE.

## 1 Background

Epilepsy is one of the most common neurological disorders and is characterized by spontaneous recurrent seizures (SRSs) [1]. Globally, over 70 million people have been diagnosed with epilepsy, and the high recurrence rate of seizures can reach 83.6% [2, 3], which can impose a significant economic burden on society and health care systems. Therefore, the etiology and pathophysiological processes of epilepsy have been largely explored, and antiepileptic drugs have been developed [4]. However, approximately one-third of epilepsy patients remain unresponsive to medication and are classified as patients with drug-resistant epilepsy. The main reason is that its etiology and pathological mechanisms are complex and not yet fully understood [5, 6]. Temporal lobe epilepsy (TLE), one of the most common types of drug-resistant epilepsy, requires further investigation into its neuropathological mechanisms, which could provide experimental evidence for identifying specific targets for the prevention and treatment of drug-resistant epilepsy.

TLE is caused by the hypersynchronous discharge of neurons in the brain resulting from an imbalance between neuronal excitation and inhibition [7]. Thus, neurons in a specific brain region become hyperexcited and their signals spread across brain areas through abnormal neural circuits, leading to hyperexcitation and abnormal synchronous firing of neurons across the global cerebral cortex, ultimately resulting in SRSs [8, 9]. Since rapid depolarization mediated by alpha-amino-3-hydroxy-5-methyl-4 isoxazole receptors (AMPARs) is crucial for the initial synchronization of synaptic transmission in the generation of synchronous network activities [10], the abnormal activation and function of AMPARs have been indicated to be among the main driving factors of epileptic seizures. Additionally, studies have shown that AMPAR antagonists have an advantage in terminating abnormal network synchronization activities during epileptic states [11–14]. Therefore, developing antagonists that target AMPARs could become a major research field for clinical antiepileptic treatments [14–16].

Functional AMPARs are assembled from four core subunits, GluA1, GluA2, GluA3 and GluA4, together with various interacting auxiliary subunits [17, 18]. Recent studies have shown that the auxiliary subunits of AMPARs can regulate AMPAR assembly, trafficking, gating, and pharmacology [19–21], which ultimately determines the composition and function of AMPARs [22–24]. These findings indicate that exploring the regulatory role of AMPAR auxiliary subunits in the function of this receptor is crucial and essential for developing treatment methods for TLE [25]. Cystine-knot AMPAR-modulating protein 44 (CKAMP44) has been identified as a novel auxiliary subunit of AMPAR and has been shown to regulate AMPAR function and synaptic plasticity [26, 27]. CKAMP44 is significantly upregulated in almost all major neuronal layers in epilepsy patients, according to a single-cell transcriptomic analysis of more than 110,000 neurons in cortical samples from the temporal lobe of epilepsy patients and nonepileptic subjects, suggesting that CKAMP44 may be involved in the pathological processes of epilepsy [28]. However, its role in TLE is not yet clear and requires further exploration.

Therefore, we hypothesize that CKAMP44 primarily participates in the pathological process of TLE by affecting AMPAR-mediated synaptic transmission and related molecular mechanisms. Our results indicated that the inhibition of CKAMP44 could reduce the susceptibility to and severity of epileptic seizures in kainic acid (KA)-induced TLE mice. Through a transcriptomic analysis and rescue experiments, we explored the mechanism of CKAMP44 in TLE. The results indicated that the inhibition of CKAMP44 downregulated the mRNA and protein levels of protein phosphatase 3 regulatory subunit B (PPP3r2), resulting in increased phosphorylation of GluA1-Ser831 and increased membrane levels of the GluA1 protein. This study validated the role of the CKAMP44-PPP3r2-GluA1 pathway in the pathological mechanism of TLE. These findings could provide a new potential target for the development of clinical antiepileptic treatments.

## 2. MATERIALS AND METHODS

### 2.1 Animals

In this study, 6-week-old C57BL/6J adult male mice weighing 20–25 g were used. The mice were subjected to standard laboratory conditions at 22 ± 1 °C with 50%–60% humidity on a 12:12-h light‒12-h dark cycle and provided plenty of water and food. All experimental animals used in our study were acquired from Chongqing Medical University Experimental Animal Center. All animal experiments were approved by the Ethics Committee of Chongqing Medical University. All efforts were made to minimize animal suffering and the number of animals included in the experiments.

### 2.2 KA-induced TLE mouse model

The mice were anesthetized and placed in a stereotaxic apparatus (RWD Life Science Co. Ltd., China). Furthermore, 0.5 μl of KA (0.3 μg in 0.5 μl of saline) was injected into the right hippocampal CA1 region for 3 minutes. The mice were allowed to recover from anesthesia at 32 °C and then monitored closely during SE. SE usually began 15 minutes after complete recovery and ended 40 minutes after SE onset with an intraperitoneal injection of diazepam (10 mg/kg). Sham control mice were treated identically but received the same volume of saline. The behavioral seizure scores were classified according to Racine’s scale [29]. Mice with seizures (Racine’s stages VI-V) observed by video monitoring were considered successful KA-induced TLE models, and the success rate of KA modeling was 100%.

### 2.3 Video monitoring and local field potential (LFP) recording

#### 2.3.1 Video monitoring

For the behavioral tests, video monitoring started when KA was injected at 4 weeks after the AAV injection. Mice were video monitored to quantify the latency of the first SRS and the number of SRSs recorded over 30 consecutive days after the KA injection. Nonconvulsive status epilepticus was terminated with diazepam two hours after the KA injection. Seizures (Racine’s stages VI-V) recorded in the last 7 days (21^st^ day to 28^th^ day) of one month with the continuous video SRS recording system were quantified during offline video review by two independent observers who were blinded to the treatment.

#### 2.3.2 LFP recordings

Surgery was performed 1 week prior to the recordings. Briefly, the mice were anesthetized and placed in a stereotaxic apparatus, and two stainless steel screws were implanted in the anterior cranium to serve as the ground electrode when recording the hippocampal LFPs. For LFP recordings, the recording electrode was implanted into the right dorsal hippocampus and connected to the MAP data acquisition system (Plexon; Texas, USA) to collect data. A typical seizure-like event was characterized by the disappearance of a normal electrophysiological rhythm and the occurrence of a cluster of spontaneous paroxysmal discharges with amplitudes greater than twice the baseline, a frequency greater than 3 Hz, and a duration greater than 5 seconds.

### 2.4 Promoter luciferase reporter assay

The promoter region of the CKAMP44 gene, extending from the −2000 base site to the transcription start site, was integrated into the modified pGL3-basic vector (GeneCreate, Wuhan, China). The full-length sequence of Nkx2-1 was extracted from the cDNA of BV2 cells and cloned and inserted into the pcDNA3.1 vector. For plasmid transfection, HEK293T cells were cultured on 24-well plates and transfected with the indicated plasmids 48 h before harvest. The activity of firefly luciferase and the signal of Renilla luciferase were quantified using a dual-luciferase reporter assay kit (GeneCreate, Wuhan, China) and detected using a GloMax 20/20 Luminometer.

### 2.5 Immunostaining

For immunofluorescence staining of cryostat sections, the mice were anesthetized and perfused with phosphate-buffered saline (PBS) followed by 4% paraformaldehyde (PFA). The brains were subsequently fixed with 4% PFA overnight and submerged in 30% sucrose for 3 days. Brain sections (10 μm) were sliced using a Leica freezing microtome. The brain sections were permeabilized with 0.4% Triton X-100 for 20 min, blocked with the goat serum working solution (Boster Biological Technology; Wuhan, China) for 60 min at room temperature, and then incubated with mixed primary antibodies diluted in PBS at 4 °C overnight. The sections were washed three times with PBS and incubated with appropriate fluorescent dye-conjugated secondary antibodies for 60 min at room temperature. 4’,6-Diamidino-2-phenylindole (DAPI) was used for nuclear staining. For immunocytochemical assays, primary neurons were fixed with a PBS solution containing 4% PFA and 4% sucrose (30 min at room temperature). The neurons were permeabilized with 0.1% Triton X-100 for 10 min and blocked with the goat serum working solution for 60 min at room temperature. Then, the neurons were incubated with mixed primary antibodies diluted in PBS at 4 °C overnight. Next, the neurons were washed three times with PBS and incubated with fluorescent dye-conjugated secondary antibodies at room temperature for 1 h. All the images were acquired under a confocal microscope (Leica; Wetzlar, Germany).

### 2.6 Antibodies and reagents

The antibodies used in this study are described in **Appendix A**. HiScript II Q RT SuperMix for qPCR (+gDNA wiper) and ChamQ Universal SYBR qPCR Master Mix were purchased from Vazyme Biotech Co., Ltd. (Nanjing, China). Protein A/G magnetic beads, chloroquine, and Ro 25-6981 were purchased from MedChemExpress (New Jersey, USA). Plasma Membrane Protein Isolation kits were purchased from Invent Biotechnologies (Minnesota, USA). Tetrodotoxin was obtained from Tocris Bioscience (Bristol, UK). Other reagents not mentioned here were purchased from Sigma‒Aldrich (St. Louis, USA).

The adeno-associated virus (AAV) encoding an siRNA targeting the full-length mouse Shisa9 sequence (shisa9-siRNA, vector: GV478-U6-MCS-CAG-EGFP) and the control virus (AAV-Control, vector: CON305-U6-MCS-CAG-EGFP) were purchased from Genechem Corp., Ltd. (Shanghai, China). AAV-PPP3r2 encoding the full-length mouse PPP3r2 sequence (vector: pAAV-CMV-PPP3r2-3xFLAG-P2A-mCherry-WPRE) and control virus (vector: pAAV-CMV-MCS-3xFLAG-P2A-mCherry-WPRE) were purchased from Obio Technology Co., Ltd. (Shanghai, China).

### 2.7 Western blot

As hippocampal function plays a crucial role in the pathological mechanism of TLE in mice, total proteins and plasma membrane proteins were extracted from the right hippocampus of the mice 28 days after the KA injection. The extraction steps were performed according to the manufacturer’s instructions. Total protein was extracted from the tissues with RIPA lysis buffer (Beyotime; Shanghai, China). Lysates containing equal amounts of protein were resolved via sodium dodecyl sulfate‒polyacrylamide gel electrophoresis (SDS‒PAGE) and transferred to polyvinylidene difluoride membranes (Millipore, USA). The PVDF membranes were incubated with 1x protein-free rapid blocking buffer (Epizyme Biotech, Shanghai, China) at room temperature for 20 minutes and then incubated with primary antibodies diluted in primary antibody dilution buffer (Beyotime) at 4 °C overnight. After 12–16 h, the PVDF membranes were washed with TBST and subsequently incubated with horseradish peroxidase (HRP)-conjugated secondary antibodies diluted in TBST at room temperature for 1 hour. The signal was detected using an enhanced chemiluminescence reagent (Biosharp, Guangzhou, China) and a Fusion FX7 image analysis system (Vilber Lourmat, Marne-laVallée, France). Blotting was quantified using ImageJ software. The target protein bands were normalized to the corresponding glyceraldehyde-3-phosphate dehydrogenase (GAPDH) or calnexin reference bands for quantification, and then the experimental group was normalized to the control group.

### 2.8 Immunoprecipitation

Hippocampal tissue samples were lysed with RIPA lysis buffer (Beyotime) supplemented with a protease inhibitor cocktail (MedChemExpress) at 4 °C for 30 minutes. Aliquots of the protein extracts were retained as input samples. Lysates containing equal amounts of proteins were incubated with Protein A/G magnetic beads (MedChemExpress) bound to anti-CKAMP44 or anti-GluA1 antibodies at 4 °C for 4 hours, according to the manufacturer’s instructions. The beads were then washed 3 times with TBST buffer. The bound proteins were eluted in protein loading buffer at 95 ℃ for 5 minutes. The samples were then processed for Western blot analysis in accordance with the experimental procedures detailed above.

### 2.9 RNA sequencing (RNA-Seq)

The RNA extraction, library construction, sequencing and bioinformatics analysis of the RNA-Seq data were conducted by Zhongke New Life Biotechnology Co., Ltd., Shanghai, China. The right hippocampi of the siCtrl-KA and siCKAMP44-KA groups (n=7 mice/group) were collected for further analysis. RNA isolation, quality control, library construction and sequencing were performed by Zhongke New Life Biotechnology. The library preparations were then subjected to sequencing on the Illumina NovaSeq 6000 platform. Differential expression analyses were conducted using Metascape, with a cutoff set at a 2-fold change and a significance threshold of P < 0.05. The pathway enrichment analysis was performed using the Kobas cluster algorithm (http://bioinfo.org/kobas) to examine the enriched KEGG pathways. Plots were generated using https://www.omicshare.com/tools/Home/Soft/heatmap.

### 2.10 Real-time quantitative PCR

After total RNA was extracted from the hippocampi of the mice using RNAiso plus reagent (Vazyme, Nanjing, China) according to the manufacturer’s instructions, cDNA was obtained using the PrimeScript^TM^ RT Reagent Kit with gDNA Eraser (Vazyme, Nanjing, China). Subsequently, real-time quantitative reverse transcription polymerase chain reaction (RT‒qPCR) was conducted with SYBR Premix Ex Taq II (Vazyme, Nanjing, China) to measure the levels of the target mRNAs using a 7900HT Fast Real-Time PCR system (Applied Biosystems). Gapdh was utilized as the internal control. Each reaction was performed in triplicate. The values were normalized to Gapdh expression to calculate the relative RNA expression levels.

### 2.11 Whole-cell patch-clamp recordings

#### 2.11.1 Slice preparation

Acute hippocampal slices were prepared from mice that were transfected with Ctrl-siRNA or CKAMP44-siRNA for 4 weeks. The animals were anesthetized with pentobarbital and transcardially perfused with ice-cold oxygenated (95% O_2_, 5% CO_2_) slicing solution (2.5 mM KCl, 1.25 mM NaH_2_PO_4_, 6 mM MgCl_2_, 1 mM CaCl_2_, 220 mM sucrose, 26 mM NaHCO_3_, and 10 mM D-glucose). After decapitation, the brains were removed for sectioning in the same ice-cold slice solution using a vibratome (Leica VT1200S, Germany). For whole-cell patch-clamp recordings, 300 μm coronal hippocampal sections were prepared. Slices were recovered in oxygenated artificial cerebrospinal fluid (aCSF; 124 mM NaCl, 3 mM KCl, 1.23 mM NaH_2_PO_4_, 26 mM NaHCO_3_, 2 mM MgCl_2_, 2 mM CaCl_2_, and 10 mM D-glucose, with the pH adjusted to 7.4) at 32 °C for 30 min and at room temperature (25 °C) for an additional 30 min before recording.

#### 2.11.2 Whole-cell patch-clamp recordings

A coronal slice containing the hippocampus was placed in a submersion chamber maintained at room temperature and perfused at 3 ml/minute with oxygenated Mg^2+^-free aCSF. All recordings were performed with patch pipettes with a resistance between 3 and 6 MΩ. We performed recordings in the whole-cell configuration using a Multiclamp 700B amplifier (Axon, USA) with 10 kHz digitization and a 2 kHz low-pass Bessel filter. Data were acquired and analyzed using pCLAMP 11.1 software (Molecular Devices Co.; San Jose, CA, USA). Series resistance changes were monitored throughout the experiment, and neurons were discarded if the series resistance surpassed 25 MΩ or changed by >20%. Neurons with unstable resting potentials or potentials >−50 mV were discarded. Transfected hippocampal CA1 pyramidal neurons were identified via eGFP fluorescence and a pyramidal somatic shape. For current-clamp recordings, the internal mixture contained 60 mM K_2_SO4, 40 mM HEPES, 60 mM NMG, 4 mM MgCl_2_, 0.5 mM BAPTA, 12 mM phosphocreatine, 2 mM Na_2_ATP, and 0.2 mM Na_3_GTP (pH 7.3, 275–290 mOsm). D-APV (50 μM), DNQX (20 μM) and picrotoxin (100 μM) were added to block synaptic transmission. A current step protocol was used to evoke action potentials by injecting 500 ms long depolarizing current steps of increasing amplitude from –100 pA to 300 pA (△20 pA).

For voltage-clamp recordings, the internal solution used to record spontaneous excitatory postsynaptic currents (sEPSCs) contained 130 mM Cs-methanesulfonate, 10 mM HEPES, 10 mM CsCl, 4 mM NaCl, 1 mM MgCl_2_, 1 mM EGTA, 5 mM NMG, 5 mM MgATP, 0.5 mM Na_3_GTP, and 12 mM phosphocreatine (pH 7.25; 280–300 mOsm). The sIPSCs were recorded using internal solutions that contained 100 mM CsCl, 10 mM HEPES, 1 mM MgCl_2_, 1 mM EGTA, 30 mM NMG, 5 mM MgATP, 0.5 mM Na_3_GTP, and 12 mM phosphocreatine (pH 7.25; 280–300 mOsm). The cells were clamped at −70 mV throughout the experiment, unless stated otherwise. Recordings were performed in the presence of 100 μM picrotoxin for sEPSCs, 20 μM DNQX and 50 μM D-APV for spontaneous inhibitory postsynaptic currents (sIPSCs), and 100 μM PTX and 100 μM DL-AP5 for AMPAR-eEPSCs. The amplitudes and frequencies of sIPSCs and sEPSCs were analyzed using MiniAnalysis software (Synaptosoft).

### 2.12 Human cerebral tissue

In this study, patients were diagnosed with refractory TLE based on the criteria established by the International League Against Epilepsy [1]. Six temporal samples of the neocortex were acquired from patients with refractory TLE as the epilepsy group. Samples were obtained from 6 patients with severe traumatic brain injury (TBI) as the control group for the brain tissue analysis. All cerebral tissues used in this study were collected from patients at the First Affiliated Hospital of Chongqing Medical University. Written informed consent was obtained from patients for the use of their brain tissue and access to their medical records for research purposes. The collection and use of all the samples were approved by the Ethics Committee of The First Affiliated Hospital of Chongqing Medical University and were conducted in accordance with the Declaration of Helsinki. The brain tissue samples obtained during the operation were immediately stored in liquid nitrogen until use. The clinical characteristics of the patients, such as age, sex, course and ASMs used before surgery, are presented in **Appendix Table 1**.

### 2.13 Statistics

GraphPad Prism 8 software (San Diego, CA, USA) and IBM SPSS Statistics 23 (New York, USA) were used for statistical analyses. The experimental data are presented as either medians with interquartile ranges or means ± standard deviations (SDs). p < 0.05 (two-tailed) was considered statistically significant. The normality of the data was analyzed using the Shapiro‒Wilk test. When the variance of the dataset was significantly different, we performed a nonparametric statistical analysis. Two-tailed Student’s t tests were used to compare two independent groups. For multiple group comparisons, one-way analysis of variance (ANOVA) was employed where appropriate. The statistical methods used for each experiment are detailed in the corresponding figure legend.

## 3. Results

### 3.1 CKAMP44 expression in TLE patients and KA-induced TLE mice

In this study, 6 TLE patients and 6 TBI patients were included, and no significant differences in age or sex were observed between the two groups (P>0.05), indicating that the human samples were comparable **(Appendix Table 2**). The expression of the CKAMP44 protein in the brain tissues of TLE patients and TBI patients was examined using Western blotting (WB) to clarify the potential role of CKAMP44 in TLE.WCompared with that in the control group (TBI patients), CKAMP44 protein expression was significantly increased in TLE patients **(Figure 1A).** Furthermore, KA-induced TLE model mice were generated, and the expression of the CKAMP44 protein in the hippocampal tissues of KA-induced TLE model mice was detected using WB on the 28th day. Compared with the control group, the expression of CKAMP44 was significantly increased in the hippocampal tissues of TLE mice **(Figure 1B)**. In addition, the localization of CKAMP44 was further detected in the CA1 and CA3 regions of the hippocampus in KA-induced TLE mice using immunofluorescence staining. The results revealed that CKAMP44 was primarily colocalized with NeuN (a neuronal marker) **(Figure 1C, E)**, whereas CKAMP44 was not colocalized with GFAP (an astrocyte marker) **(Figure 1D, F)**. These findings indicated that CKAMP44 expression increased significantly and was primarily localized in neurons in the hippocampal tissues of KA-induced TLE mice, suggesting that CKAMP44 may play a role in TLE by directly regulating neuronal function.

**Figure 1.**
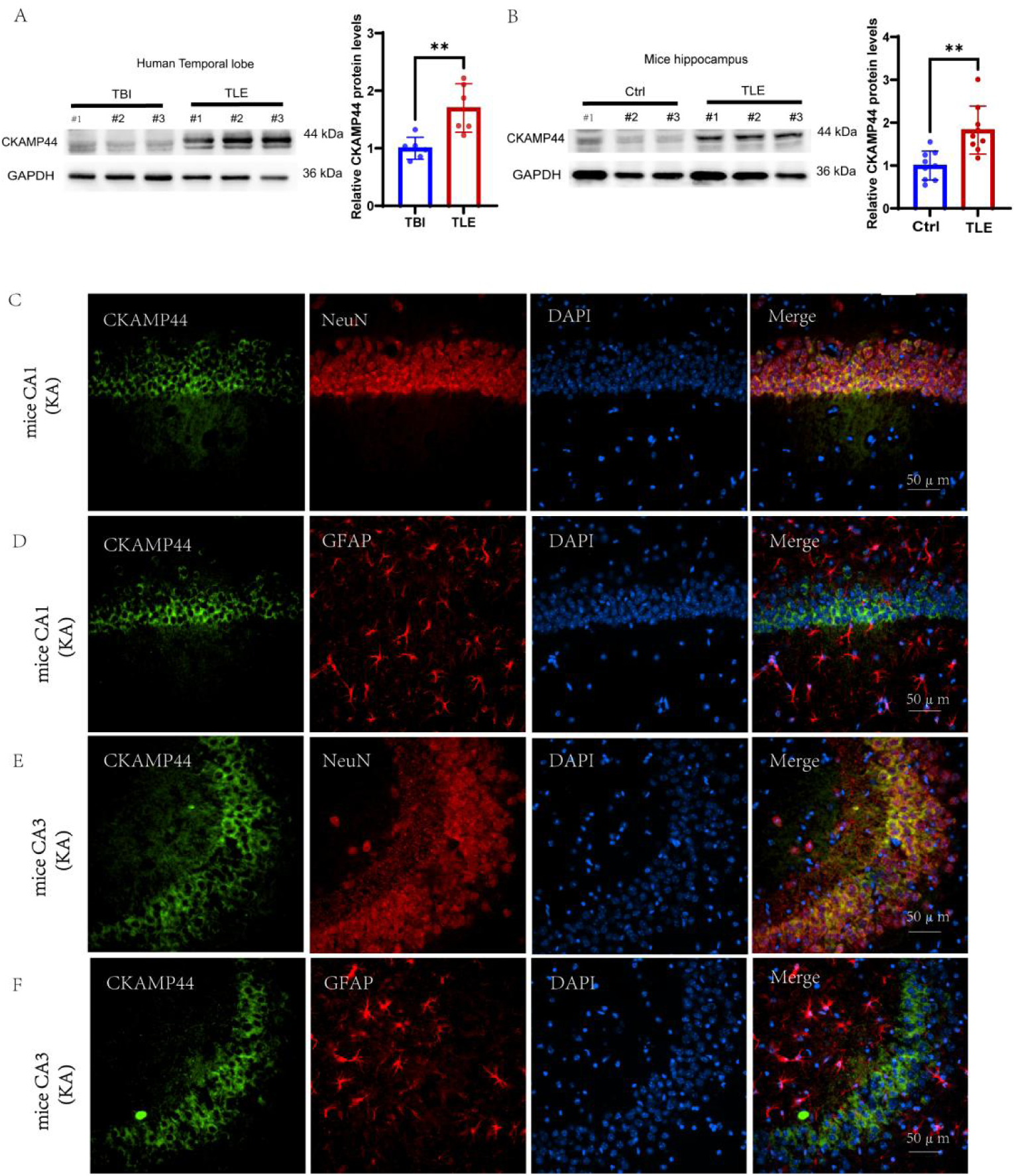
The expression and location of CKAMP44 in the brain tissues of TLE patients and KA-induced TLE mice. A. Representative WB and statistical graphs of CKAMP44 protein expression was detected in TLE patients. n=6/group; \*\**P*<0.01. B. Representative WB and statistical graphs of CKAMP44 protein expression were detected in KA-induced TLE mice. n=9/group; \*\**P*<0.01. C. CKAMP44 was primarily co-localized with NeuN (the neuronal marker) in the hippocampus CA1 of KA-induced TLE mice. D. CKAMP44 was not co-localization with GFAP (the astrocyte marker) in the hippocampus CA1 of KA-induced TLE mice. E. CKAMP44 was primarily co-localized with NeuN (the neuronal marker) in the hippocampus CA3 of KA-induced TLE mice. F. CKAMP44 was not co-localized with GFAP (the astrocyte marker) in the hippocampus CA3 of KA-induced TLE mice.

### 3.2 Inhibition of CKAMP44 suppressed seizure activity in KA-induced TLE mice

Since CKAMP44 expression was abnormally elevated during the chronic phase of TLE, we hypothesized that CKAMP44 knockdown could suppress seizure activity. First, CKAMP44-siRNA was stereotactically injected into the CA1 region of the hippocampus, and after 28 days, the infection efficiency of siCKAMP44-siRNA was validated using immunofluorescence staining and WB. Immunofluorescence experiments revealed that the hippocampus was successfully infected with the virus by the autofluorescence of AAV-eGFP in the hippocampal region **(Figure 2A)**. The efficiency of virus transfection was further detected by WB. Compared with the siCtrl group, the expression of the CKAMP44 protein was significantly decreased in the hippocampal tissue of the siCKAMP44 group **(Figure 2B-C)**. These results showed that the local stereotactic injection of CKMAP44-siRNA into the hippocampus effectively reduced the expression level of the CKAMP44 protein.

**Figure 2.**
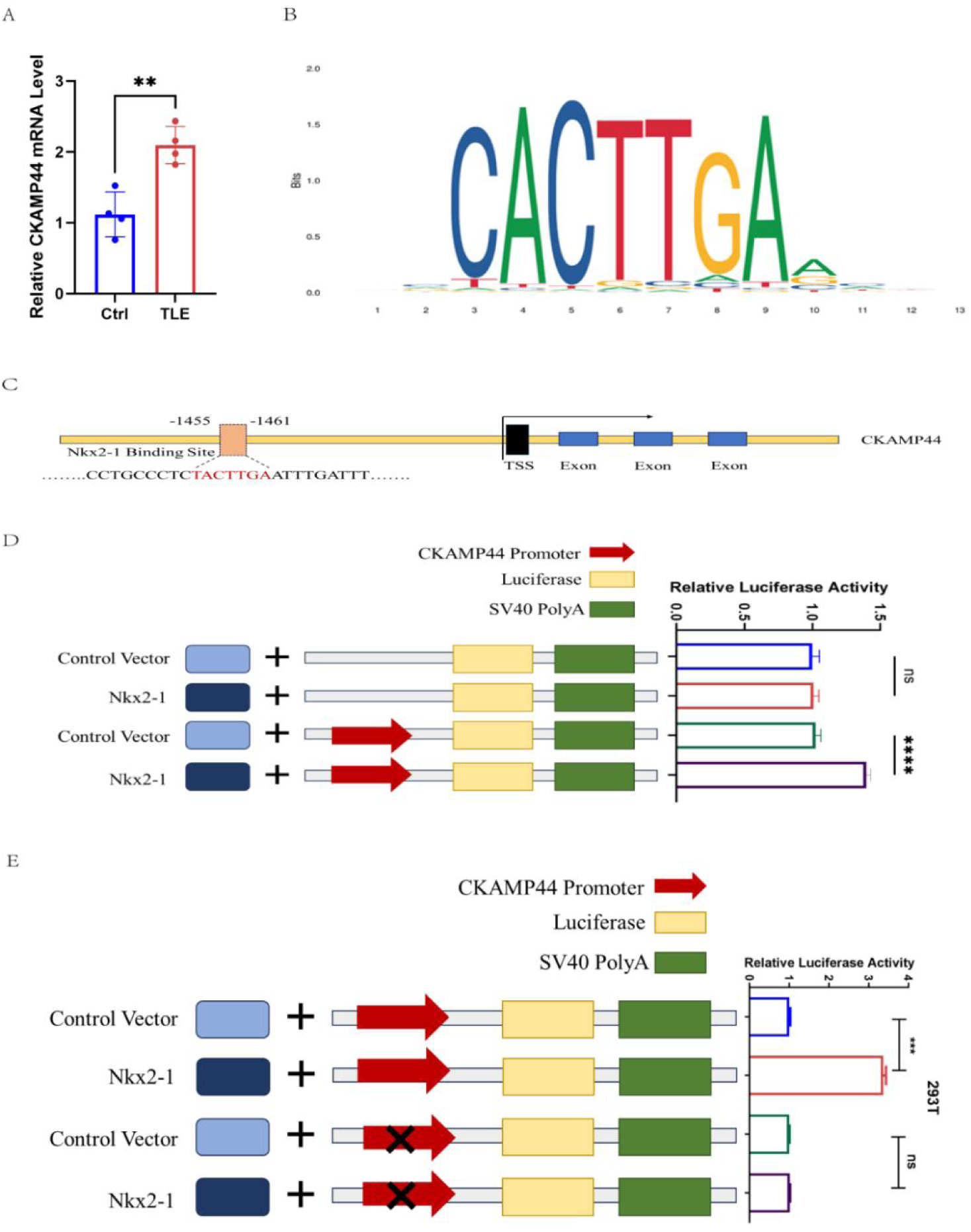
Nkx2-1 regulates the transcriptional levels of CKAMP44 mRNA. A. Compared to the control group, the expression of CKAMP44 mRNA increased significantly in the hippocampal tissues of TLE mice. n=4/group; \*\**P*<0.01. B. The potential binding site of Nkx2-1 to the CKAMP44 promoter region. C. The presence of a potential binding site upstream of the CKAMP44 transcription start site. D. The dual-luciferase reporter assay results indicated that Nkx2-1 strongly promoted the transcriptional activity of the CKAMP44 gene. E. The dual-luciferase experiment indicated Nkx2-1 could not promote the transcriptional activity of CKAMP44 by mutating the 1455 to 1461 promoter region of CKAMP44.\*\**P* < 0.01, \*\*\*\**p*<0.001; ns, no significant significance; unpaired two-tailed Student’s t-test.

Then, KA was injected on the 28^th^ day after the CKAMP44-siRNA injection, and the behaviors of the mice were continuously monitored to evaluate their seizure susceptibility and the seizure severity on the 28^th^ day after the KA injection **(Figure 2D)**. Compared with the siCtrl-KA group, the siCKAMP44-KA group presented a prolonged latency to the first onset of SRSs **(Figure 2E)** and a reduced number of grade VI-V SRSs during the 21^st–^28^th^ days **(Figure 2F)**. Additionally, the hippocampal LFP recordings revealed fewer seizures and a shorter duration of SLE in the siCKAMP44-KA group than in the siCtrl-KA group **(Figure 2G-J)**. These results indicated that the inhibition of CKAMP44 suppressed seizure activity in KA-induced TLE mice.

### 3.3 Nkx2-1 promoted the transcription of the CKAMP44 mRNA

We further verified the expression of the CKAMP44 mRNA in the hippocampal tissues of KA-induced TLE mice on the 28th day via qPCR. Compared with the control group, the expression of the CKAMP44 mRNA increased significantly in the hippocampal tissues of TLE mice **(Figure 3A)**. Next, we explored the possible regulatory mechanisms upstream of CKAMP44 according to the transcriptomic results of hippocampal brain tissues from mice in the siCtrl-KA group and the siCKAMP44-KA group **(Appendix Figure 1)**. Nkx2-1 has been reported to be involved in the pathological mechanism of epilepsy [30]. Next, we asked whether Nkx2-1 regulates the transcription of CKAMP44. We addressed this question by predicting the potential binding site of Nkx2-1 to the CKAMP44 promoter region using the JASPAR website. The analysis revealed the presence of a potential binding site upstream of the CKAMP44 transcription start site **(Figure 3B-C)**. Furthermore, we conducted a dual-luciferase reporter experiment to determine whether Nkx-2-1 promotes the transcription of CKAMP44. The dual-luciferase reporter assay results indicated that Nkx2-1 strongly promoted the transcription of the CKAMP44 gene **(Figure 3D)**. We subsequently performed a dual-luciferase experiment to verify the interaction between Nkx2-1 and the CKAMP44 promoter region. The results of the dual-luciferase experiment indicated that Nkx2-1 could not promote the transcription of CKAMP44 when bases 1455 to 1461 in the CKAMP44 promoter region were mutated **(Figure 3E)**.

**Figure 3.**
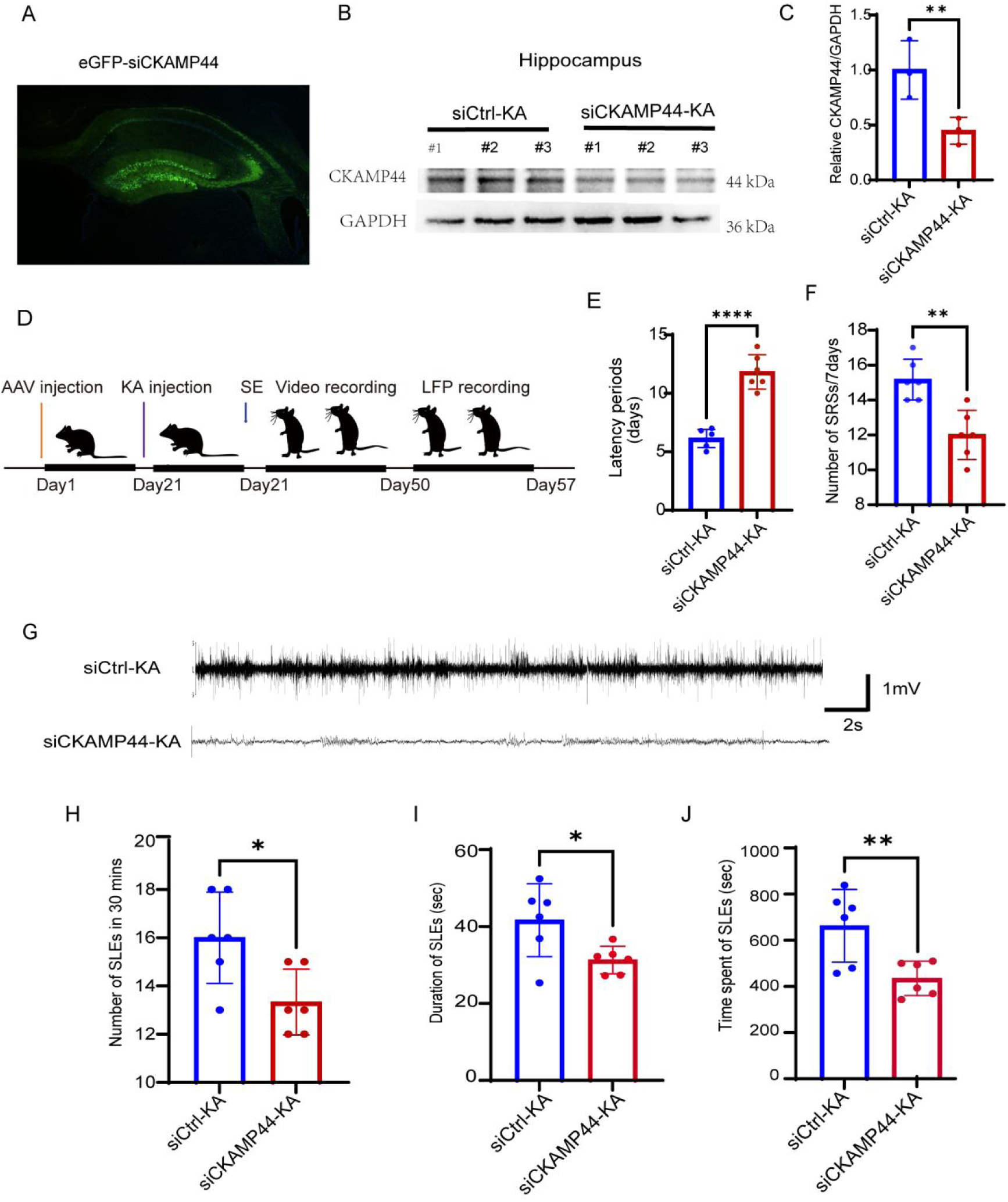
Inhibition of CKAMP44 suppressed seizure activity in KA-induced TLE mice. A. Stereotactic injection of CKAMP44-siRNA into the CA1 of the hippocampus, immuno-fluorescence is used to detect the representative expression of eGFP after 28days. B-C. Stereotactic injection of KA after 28days after CKAMP44-siRNA into the CA1 region of the hippocampus, and representative WB and statistical graphs of CKAMP44 protein expression were detected after 28days from KA injection. n=3/group; \*\**P*<0.01. D. Behavioral video monitoring and LFP recording patterns during SRS seizures in KA induced TLE mice. E-F. The effect of siCKAMP44 on the latency of SRS onset and the number of grade VI-V SRS recorded during 21-28 days (7days) recording period from the time of KA injection. n = 6/group; \*\**p*<0.01; \*\*\*\**p*<0.001; unpaired two-tailed Student’s t-test. G. Typical LFP recordings during SRS seizures in KA induced TLE mice. H-J. The effect of siCKAMP44 on the incidence and duration of SLE in KA induced TLE mice. n = 6/group; \**p*<0.05; \*\**p*<0.01; unpaired two-tailed Student’s t-test.

### 3.4 Inhibition of CAMP44 decreased glutamatergic synaptic transmission

Seizure activity occurs due to abnormal changes in the intrinsic excitability of neurons and/or synaptic transmission [31]. We first examined whether the inhibition of CKAMP44 influenced the intrinsic excitability of hippocampal CA1 pyramidal neurons by performing whole-cell patch-clamp recordings in an in vitro epilepsy model to determine whether the inhibition of CKAMP44 affects the intrinsic excitability of neurons and/or synaptic transmission. Since evoked action potentials (eAPs) reflect the level of intrinsic neuronal excitability, we first investigated the effect of CKAMP44 knockdown on eAPs in CA1 pyramidal neurons in the epilepsy model in vitro (**Figure 4A**). The results revealed no difference in the resting membrane potential **(Figure 4B)** or discharge frequency **(Figure 4C)** of CA1 pyramidal neurons between the siCtrl group and the siCKAMP44 group. These findings suggested that the knockdown of CKAMP44 did not alter the membrane properties or intrinsic excitability of neurons.

**Figure 4.**
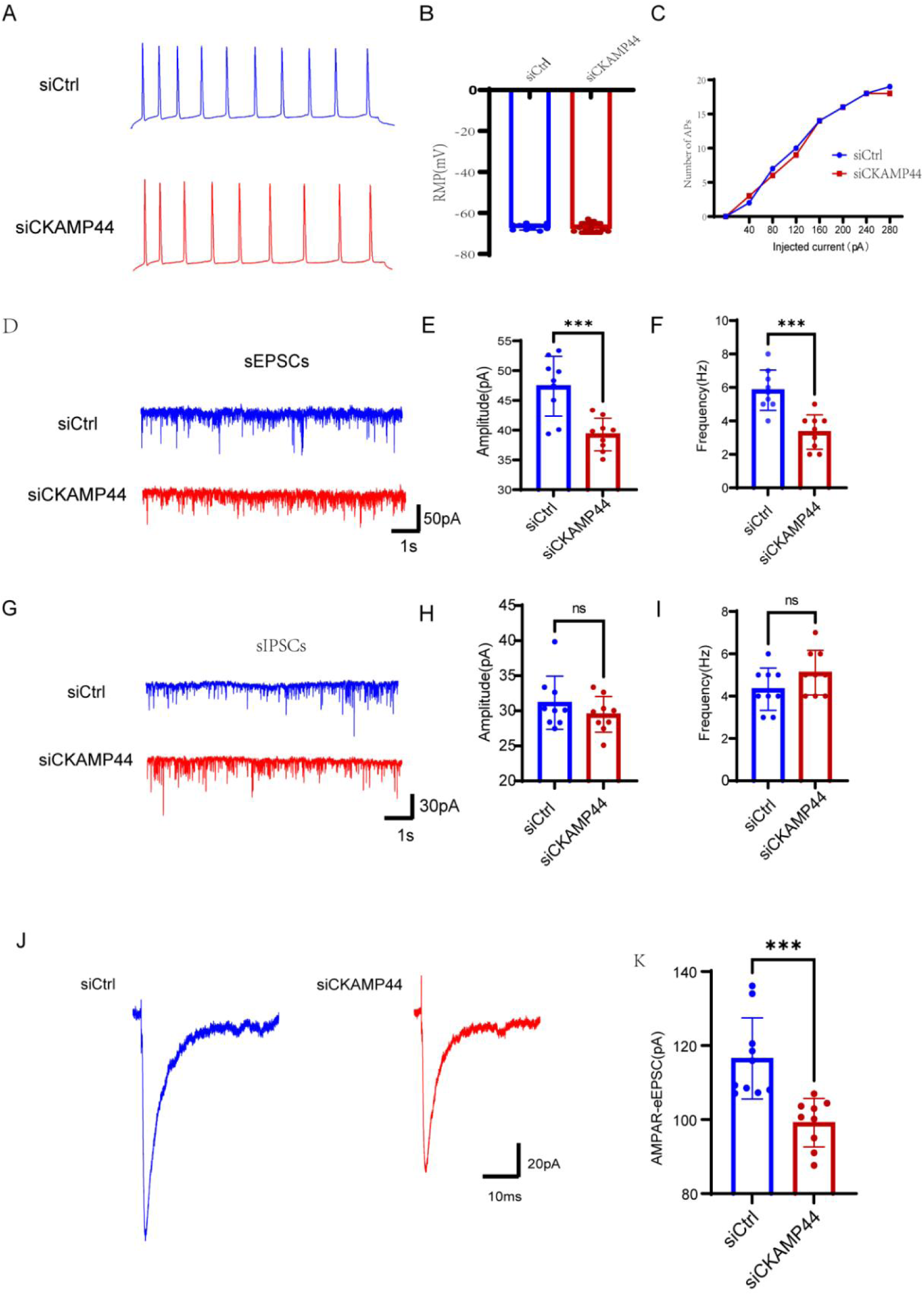
Inhibition of CKAMP44 decreased glutamatergic synaptic transmission. A. Representative graph of the eAP of CA1 pyramidal neurons in two groups. B. Statistical analyses for resting membrane potential (RMP) recorded from CA1 pyramidal neurons in two groups. n=9 cells/3 mice/group, unpaired two-tailed Student’s t-test. C. Statistical graph of recordings of the average firing rates in response to different current injections from CA1 pyramidal neurons of two groups. n = 9 cells/3 mice/group; ns, no statistical significance; two-way repeated measures ANOVA. D. Representative trajectories of the sEPSCs in CA1 pyramidal neurons in two groups. E. Statistical graph of the frequency of sEPSCs between two groups. n=9 cells/3 mice/group; \*\*\**P*<0.001; unpaired two-tailed Student’s t-test. F. Statistical graph of amplitude of sEPSCs between two groups. n=9 cells/3 mice/group. G. Representative trajectories of sIPSCs of pyramidal neurons in hippocampus CA1 in the two groups. H. Statistical graphs of the frequency of sIPSCs of pyramidal neurons in hippocampus CA1 between two groups. n=9 cells/3 mice/group; ns, no significant significance; unpaired two-tailed Student’s t-test. I. Statistical graphs of the amplitude of sIPSCs of pyramidal neurons in hippocampus CA1 between two groups. n=9 cells/3 mice/group; ns, no significant significance; unpaired two-tailed Student’s t-test. J. Representative graph of AMPAR-eEPSC of CA1 pyramidal neurons. K. Statistical graphs of the amplitude of AMPAR-eEPSC of pyramidal neurons in hippocampus CA1 between two groups. siCtrl group, n=10 cells/3 mice; siCKAMP44 group, n=9 cells/3 mice; *** *p*<0.001; unpaired two-tailed Student’s t-test.

Next, we explored whether the inhibition of CKAMP44 affected synaptic transmission in an epilepsy model in vitro. Therefore, we recorded sEPSCs and sIPSCs in the CA1 region of pyramidal neurons in the mouse hippocampus by performing whole-cell patch-clamp recordings in an in vitro epilepsy model. The results revealed that CA1 pyramidal neurons from the siCKAMP4 group presented a significantly lower frequency and lower amplitudes of sEPSCs than did those from the siCtrl group **(Figure 4D–F)**. Furthermore, no significant differences in the frequency or amplitude of sIPSCs were observed between the siCtrl group and the siCKAMP44 group **(Figure 4G-I).**

Furthermore, previous studies reported that the synchronization of hippocampal neuronal activity depends on AMPAR-mediated rapid synaptic transmission [32], and because CKAMP44 primarily affects AMPAR-mediated excitatory postsynaptic currents (AMPAR-EPSCs), we further recorded AMPAR-EPSCs in the hippocampal CA1 region of mice via whole-cell patch-clamp recordings in an epilepsy model in vitro. Compared with those in the siCtrl group, the amplitude of AMPAR-EPSCs in CA1 pyramidal neurons in the hippocampal tissues of the mice in the siCKAMP44 group was lower in the epilepsy model in vitro, suggesting that the knockdown of CKAMP44 decreased AMPAR-mediated synaptic transmission **(Figure 4J-K)**.

The above results showed that the inhibition of CKAMP44 primarily decreased the amplitude of AMPAR-EPSCs and decreased the excitatory synaptic transmission of pyramidal neurons in the hippocampal CA1 region of TLE mice.

### 3.5 Inhibition of CKAMP44 upregulated the surface expression of GluA1

The inhibition of CKAMP44 influenced the function of AMPAR, which depends on the expression of the core subunits of AMPAR; therefore, we first verified the total and surface protein expression of GluA1-GluA4 using WB. The results revealed no statistically significant changes in the total protein expression of GluA1-GluA4 in the hippocampal tissues of the mice in the siCKAMP44-KA group compared with those of the mice in the siCtrl-KA group **(Figure 5A-E)**, suggesting that CKAMP44 knockdown did not affect the total protein expression of GluA1 to GluA4.

**Figure 5.**
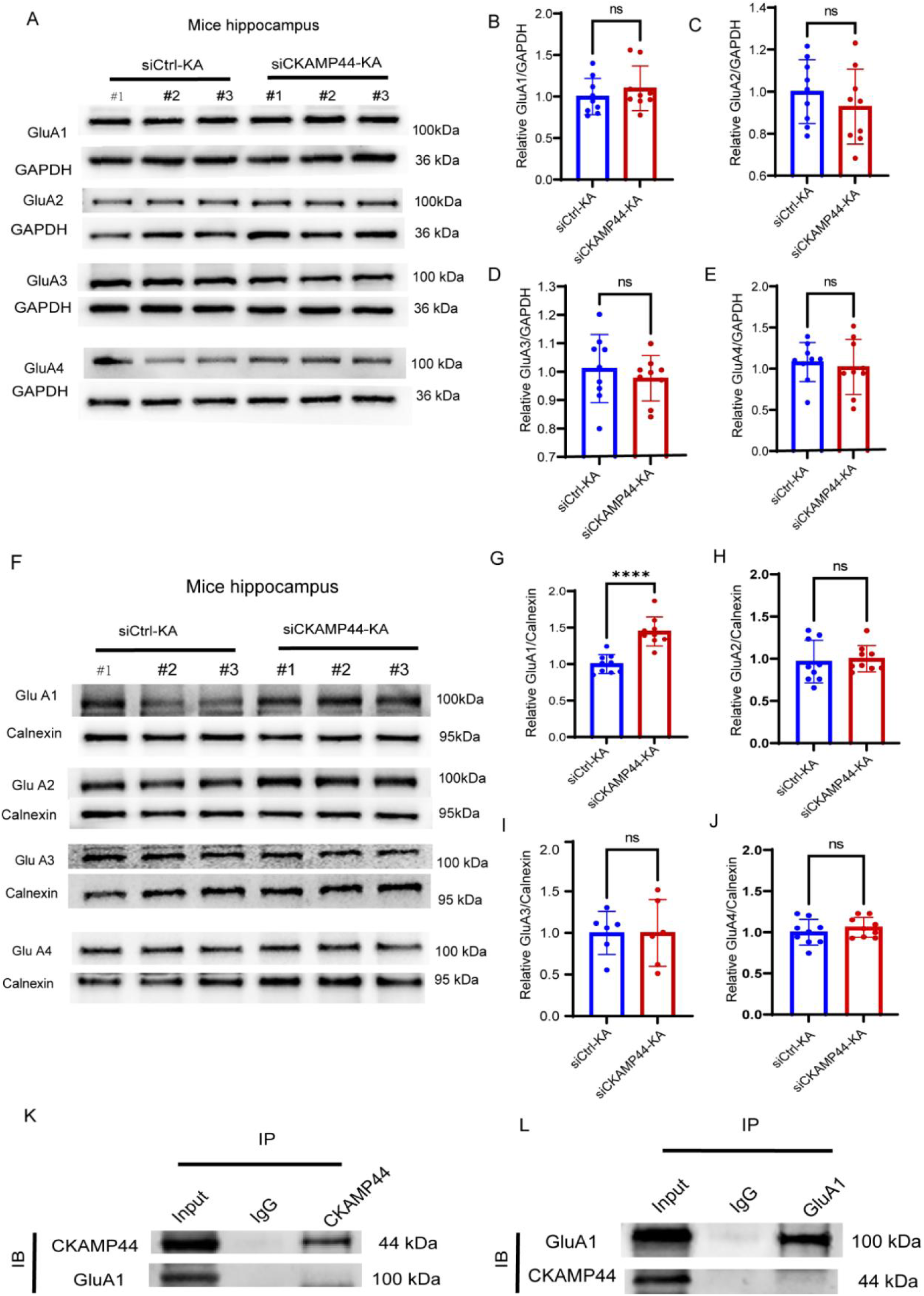
Inhibition of CKAMP44 regulated surface expression of GluA1. A. Representative WB imaging of the total protein expression of the GluA1-A4 in the hippocampal tissues of mice. B-E. Statistical graphs of the total expression of the GluA1-A4 in siCtrl-KA group and siCKAMP44-KA group. n = 9/group, ns, no significant significance; unpaired two-tailed Student’s t-test. F. Representative WB imaging of the surface expression of GluA1-A4 in the hippocampal tissues of mice. G-J. Statistical graphs of the surface expression of the GluA1-A4 in siCtrl-KA group and siCKAMP44-KA group. G-I. n = 9/group; J, n=6/group; \*\*\*\**p*<0.001; ns, no significant significance; unpaired two-tailed Student’s t-test.

Since only GluA1 to GluA4 located on the cell surface can perform synaptic transmission functions, we further verified the surface expression of GluA1 to GluA4 using WB. Compared with that in the siCtrl-KA group, the membrane surface expression of GluA1 was significantly increased in the hippocampal tissues of the siCKAMP44-KA group **(Figure 5F-G)**. However, no statistically significant differences in the surface expression of GluA2 to GluA4 were observed **(Figure 5H-J)**. These findings indicated that, in the hippocampal tissues of KA-induced TLE mice, the inhibition of CKAMP44 specifically upregulated the surface expression of GluA1. Given that CKAMP44 and GluA1 were shown to interact in previous studies, we first conducted a coimmunoprecipitation (co-IP) experiment. However, the results did not indicate an interaction between CKAMP44 and GluA1 **(Figure 5K-J)**.

### 3.6 Inhibition of CKAMP44 upregulated the surface expression of GluA1 and the phosphorylation level of GluA1-Ser831 by downregulating PPP3r2

We performed a transcriptomic analysis on the hippocampal brain tissues of the mice in the siCtrl-KA and siCKAMP44-KA groups (n=7 mice/group) to explore the possible mechanisms by which CKAMP44 affects GluA1 surface expression. The PCA results indicated that the samples from the two groups could be effectively distinguished, suggesting that the transcriptomic results were reliable **(Figure 6A**). The analysis of differentially expressed genes and enriched KEGG pathways suggested that these genes were enriched mainly in common pathways such as lipid metabolism, carbon metabolism, and glutamatergic synapses (**Figure 6C**). Considering the previous electrophysiological experimental results, CKAMP44 may participate primarily in the pathological mechanism of TLE by affecting the glutamatergic synapse pathway. Among the genes enriched in this pathway, PPP3r2 was the most significantly differentially expressed (**Figure 6B**). Therefore, we hypothesized that PPP3r2 is a key molecule in the downstream glutamatergic synapse pathway, which was validated using qPCR and WB. Both the qPCR and WB results revealed that, compared with those in the siCtrl-KA group, the mRNA and protein expression levels of PPP3r2 were significantly decreased in the hippocampal tissue of the siCKAMP44-KA group **(Figure 7A-C).**

**Figure 6.**
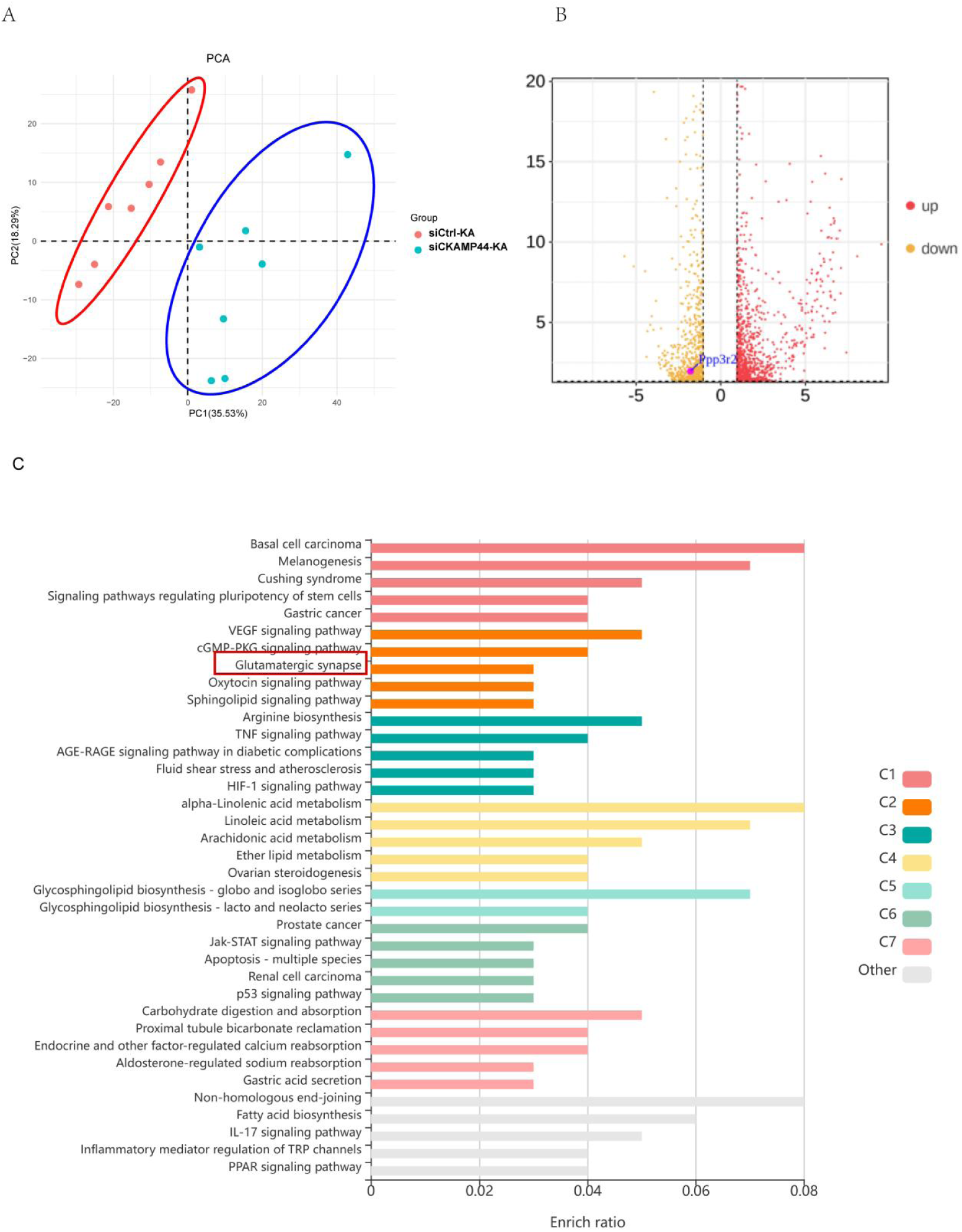
Inhibition of CKAMP44 regulated glutamatergic synaptic pathway in KA-induced TLE mice. A. The discrimination between two sample sizes, red represents the siCtrl-KA group, and green represents the siCKAMP44-KA group. B. Volcano plot of differentially expressed genes between two groups. The horizontal axis represents the value of log2FC, and the vertical axis represents the - log10 value of P value. Red represents unregulated genes, orange represents downregulated genes and blue represents total differentially expressed genes. C. KEGG enrichment analysis was performed on differentially expressed genes. Each row represents an enrichment function, and the length of the bar represents the enrichment ratio. The calculation method is “input gene count”/“background gene count”. The color of the bar is the same as the color in the circular network above, representing different clusters. n=7 mice/group.

**Figure 7.**
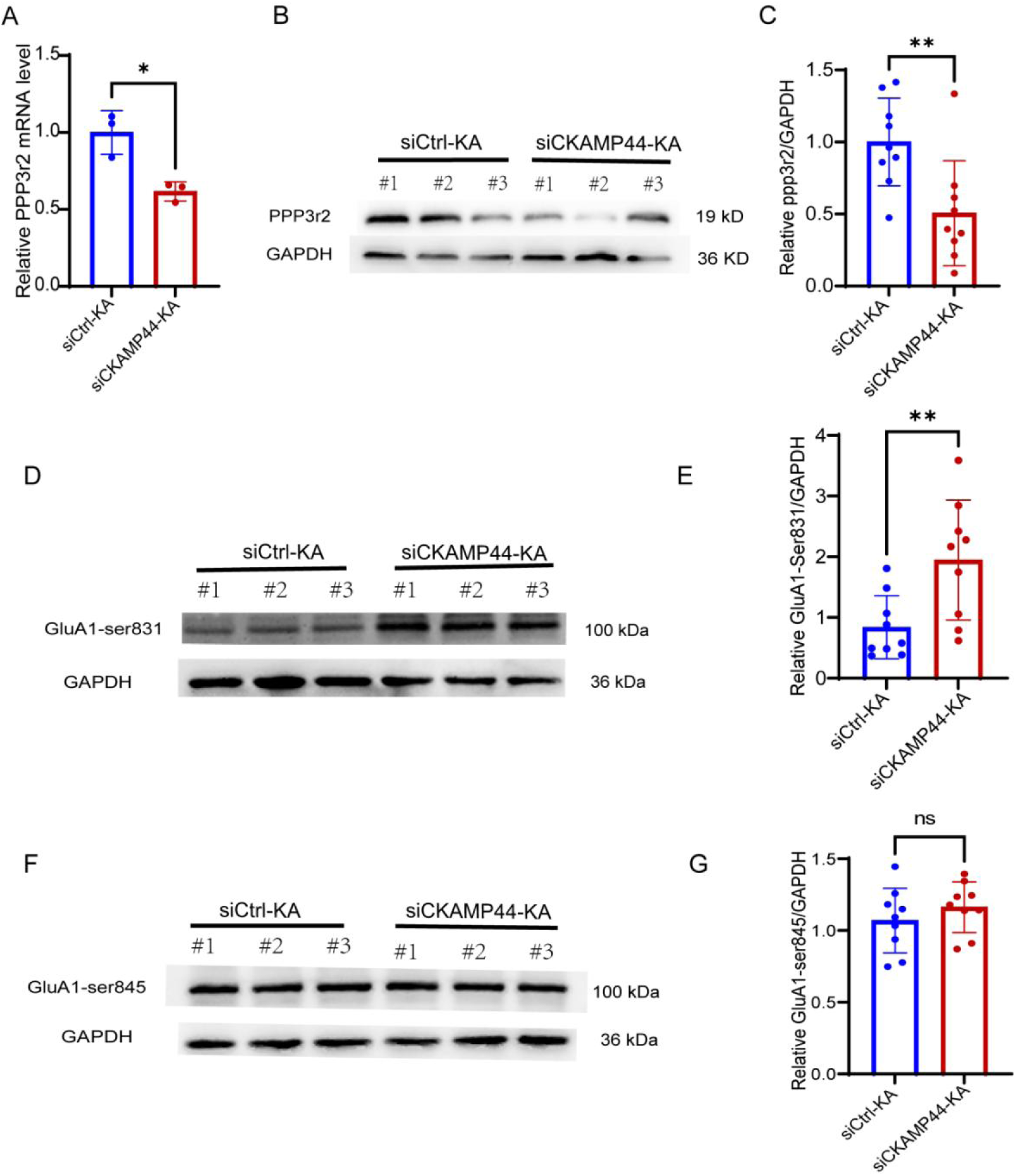
Inhibition of CKAMP44 upregulated expression of GluA1-ser831 by downregulating PPP3r2. A. Statistics of PPP3r2 mRNA in the hippocampal tissues of two groups. n=3/group; \**p*<0.05. B-C. Representative WB and statistical graphs of PPP3r2 protein expression in the hippocampus of two groups. n=9/group; \*\**p*<0.01; unpaired two-tailed Student’s t-test. D-E. Representative WB graph and statistical analysis of the phosphorylation of GluA1-Ser831. n=9/group; \*\**P*<0.01; unpaired two-tailed Student’s t-test. F-G. Representative WB graph and statistical analysis of the phosphorylation of GluA1-Ser845. n=9/group; ns, no statistical significance;unpaired two-tailed Student’s t-test.

Furthermore, PPP3r2 is a recently discovered protein phosphatase subunit that functions primarily in dephosphorylation [33]. Previous studies have reported that the surface expression and function of GluA1 are affected mainly by serine phosphorylation sites, including the GluA1-Ser831 and GluA1-Ser845 sites [34]. Therefore, their phosphorylation levels were detected via WB. Compared with that in the siCtrl-KA group, the level of phosphorylation at the GluA1-Ser831 site was significantly increased in the hippocampal tissue of the siCKAMP44-KA group (**Figure 7D-E**), whereas significant differences in the levels of phosphorylation at the GluA1-Ser845 site were not observed between the two groups (**Figure 7F-G**). Therefore, in the pathological state of TLE, the inhibition of CKAMP44 primarily increased the phosphorylation level of GluA1-Ser831 and the surface expression of GluA1 by downregulating the expression of PPP3r2.

### 3.7 Inhibition of CKAMP44 suppressed seizure activity through the CKAMP44-PPP3r2-GluA1 pathway

We further verified that the inhibition of CKAMP44 suppressed seizure activity in TLE mice primarily through the CKAMP44-PPP3r2-GluA1 pathway by simultaneously injecting the CKAMP44-siRNA and adPPP3r2-AAV into the hippocampal CA1 region of the mice. Immunofluorescence staining confirmed that siCKAMP44 and adPPP3r2-AAV successfully infected neurons in the hippocampal CA1 region of the mice, and the two viruses colocalized **(Figure 8A)**. Compared with that in the control group (siCKAMP44-adCtrl), the expression of the PPP3r2 protein was significantly increased in the hippocampal tissue of the siCKAMP44-adPPP3r2 group (siCKAMP44-adPPP3r2) **(Figure 8B)**. These results indicated that the two viruses colocalized.

**Figure 8.**
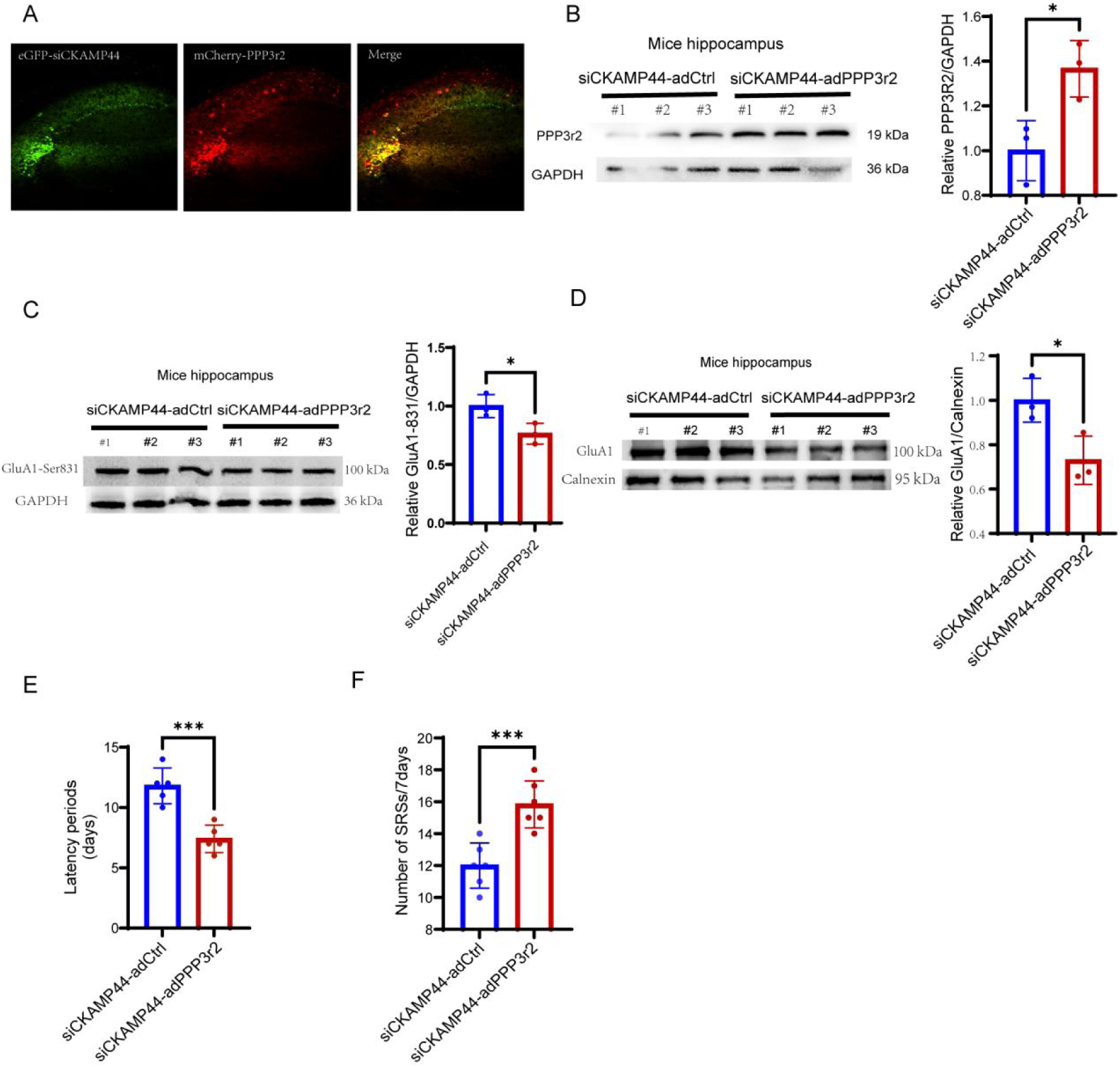
Inhibition of CKAMP44 suppressed seizure activity through CKAMP44-PPP3r2-GluA1 pathway. A. Stereotactic injection of CKAMP44-siRNA and adPPP3r2-AAV into the CA1 of the hippocampus, immunofluorescence is used to detect the representative expression of eGFP and mCherry after 28days. eGFP represented CKAMP44-siRNA; mCherry represented adPPP3r2-AAV. B. Stereotactic injection of CKAMP44-siRNA and adPPP3r2-AAV into the CA1 region of the hippocampus, and after 28days, representative WB and statistical graphs of PPP3r2 protein expression were detected. n=3/group; \**P*<0.05; unpaired two-tailed Student’s t-test. C. Representative WB and statistical graphs of GluA1-Ser831 protein expression in hippocampal tissue of KA induced TLE mice. n=3/group; *\*p*<0.05;unpaired two-tailed Student’s t-test. D. WB representative and statistical graphs of GluA1 surface protein expression in hippocampal tissue of KA induced TLE mice. n=3/group; \**p*<0.01;unpaired two-tailed Student’s t-test. E-F. KA injection was performed 28days after intervention with CKAMP44-siRNA and adPPP3r2-AAV. Statistical graphs of the latency and frequency of VI-V grade SRS in two groups within 21-28 days (7days) after KA injection. n=5/group; ** *p*<0.01; *** *p*<0.001; unpaired two-tailed Student’s t-test.

Then, the levels of GluA1-Ser831 phosphorylation and the surface expression levels of GluA1 were detected via WB, and the behavior of epileptic seizures in KA-induced TLE mice was video monitored. Compared with those in the siCKAMP44-adCtrl group, the level of GluA1-Ser831 phosphorylation **(Figure 8C)** and the surface expression of GluA1 **(Figure 8D)** were significantly decreased in the hippocampal tissue of the siCKAMP44-adPPP3r2 group. Additionally, compared with that in the siCKAMP44-adCtrl group, the latency of the first onset of SRSs was shorter, and the number of grade VI-V SRSs was greater in the KA-induced TLE mice in the siCKAMP44-adPPP3r2 group **(Figure 8E-F).** These results indicated that the overexpression of PPP3r2 could antagonize the protective effects of CKAMP44 knockdown on TLE mice. Therefore, during the chronic phase of TLE, the inhibition of CKAMP44 primarily promoted the phosphorylation of GluA1 at Ser831 and the surface expression of GluA1 by downregulating the expression of PPP3r2, ultimately suppressing seizure activity and exerting a protective effect on TLE mice.

## 4 Discussion

Several principal findings were obtained from this study. 1) The expression of the CKAMP44 protein and mRNA is significantly increased in the brain tissues of patients with TLE and KA-induced TLE mice, and Nkx2-1 regulates the transcription of CKAMP44 in the hippocampal tissues of KA-induced TLE mice. 2) The inhibition of CKAMP44 suppresses the susceptibility and severity of epileptic seizures in KA-induced TLE mice and further reduces excitatory synaptic transmission and AMPAR-mediated EPSCs in acute hippocampal slices. 3) Nkx2-1 regulates the transcription of CKAMP44 in the hippocampal brain tissues of KA-induced TLE mice;4) The inhibition of CKAMP44 suppresses seizure susceptibility and decreases seizure severity in KA-induced TLE mice by increasing the phosphorylation of the GluA1-Ser831 site and the expression level of GluA1 by downregulating the expression of PPP3r2. To our knowledge, this is the first study to demonstrate that the inhibition of CKAMP44 can suppress pathological electrical activity in the brains of TLE mice by affecting AMPAR function, which may provide a new therapeutic target for the treatment of epileptic seizures.

The KA-induced mouse model effectively simulates the pathological process of temporal lobe epilepsy [35]. In this study, the expression and localization of CKAMP44 were first examined not only in the brain tissues of TLE patients but also in those of KA-induced TLE mice. The results revealed that the expression of the CKAMP44 protein was significantly increased both in the hippocampal tissue of TLE mice and in the temporal lobe of TLE patients. We also observed that CKAMP44 mRNA levels were significantly increased in KA-induced TLE mice. This finding is consistent with previous findings that CKAMP44 mRNA expression is significantly increased in temporal lobe samples from epilepsy patients according to a single-nucleus transcriptomic analysis [28]. We performed experiments to clarify how epilepsy leads to the upregulation of CKAMP44 and found that epilepsy activated the transcription factor NKx2-1, which is involved in diverse biological processes [36]. Using a luciferase assay and reverse transcription PCR (RT–PCR) experiments, we confirmed that NKx2-1 binds to the promoter of the CKAMP44 gene and promotes its transcription. Furthermore, CKAMP44 was colocalized primarily with neurons in the CA1 and CA3 regions of the hippocampus in KA-induced TLE mice. Thus, these results indicate that the abnormally increased expression of CKAMP44 plays an important role in the pathological process during the chronic phase and that its role involves influencing the function of neurons in the hippocampus.

The epilepsy phenotype of the mice was subsequently observed through behavioral monitoring and LFP recordings, which revealed that the inhibition of CKAMP44 suppressed the susceptibility to and severity of epileptic seizures in KA-induced TLE mice. TLE is caused mainly by excessive excitation and abnormal synchronous discharge of cortical neurons. Synaptic dysfunction mediated by excitatory glutamatergic receptors plays an important role in seizures [37]. Therefore, further electrophysiological studies in acute hippocampal slices from a KA-induced mouse model revealed that the inhibition of CKAMP44 reduced excitatory glutamatergic synaptic transmission and AMPAR-mediated excitatory postsynaptic currents. This finding is consistent with previous findings that CKAMP4^−/−^ mice display a dramatic reduction in the amplitudes of extrasynaptic AMPAR-mediated currents compared with wild-type mice [38]. Since AMPAR-mediated synaptic function is related to the expression of the core subunits of GluA1 to GluA4, we found that the inhibition of CKAMP44 specifically upregulated the membrane expression of GluA1 by detecting the total protein and membrane protein expression of GluA1 to GluA4 in the hippocampal tissue of TLE mice. This finding is consistent with previous studies showing that abnormal changes in the expression and function of GluA1 are closely related to epilepsy [39, 40].

The localization, activation, and abundance of GluA1 are highly regulated [41, 42], and we further explored the underlying mechanism underlying the upregulation of GluA1. However, surprisingly, no interaction was observed between CKAMP44 and GluA1, which is different from the findings of previous studies; that is, under physiological conditions [38]. An increasing number of studies have indicated that CKAMP44 plays different roles in the pathological processes of various diseases [43–45]. Therefore, the different results of the interaction between GluA1 and CKAMP44 can be interpreted as follows: under physiological conditions, CKAMP44, a type I transmembrane protein, has a complete cysteine-rich extracellular domain and an intracellular C-terminal domain, which ensures the interaction between CKAMP44 and GluA1 and regulates its function. In particular, the extracellular domain is necessary for the physical interaction between CKAMP44 and GluA1 [38]. In the pathological state of epilepsy, the possible reason for the lack of interaction between CKAMP44 and GluA1 is that CKAMP44 may have changes in its extracellular domain, leading to its inability to interact with GluA1. Therefore, CKAMP44 may affect the membrane expression of GluA1 and AMPAR-mediated synaptic function through other mechanisms in the pathophysiological process of epilepsy, which requires further exploration.

Next, a transcriptomic analysis was performed on the hippocampi of the mice in the siCtrl-KA group and the siCKAMP44-KA group, and the results indicated that the inhibition of CKAMP44 could participate in the pathological process of TLE via the glutamatergic synaptic transmission pathway. Among all the DEGs in this pathway, PPP3r2 is considered the most critical, with the maximum fold change in expression. PPP3r2 is regulatory subunit B of protein phosphatase 3, which functions primarily in cell signaling by dephosphorylating transcription factors, ion channels, and enzymes to regulate specific molecular and cellular functions [46]. Based on the function of PPP3r2 and the altered expression of GluA1, the phosphorylation level of GluA1 was subsequently detected, and the results revealed that after the inhibition of CKAMP44, the phosphorylation level of GluA1-Ser831 increased. Therefore, the inhibition of CKAMP44 primarily increases the phosphorylation of GluA1 at Ser831 and the membrane expression level of GluA1 by downregulating the protein expression of PPP3r2. This finding is consistent with previous studies showing that phosphorylation plays a significant role in regulating the function of GluA1 and maintaining the normal electrical activity of neurons [47, 48], and dysregulation of GluA1 dephosphorylation is an important process leading to abnormal neuronal discharge and even status epilepticus [49, 50]. Therefore, in the pathological state of TLE, the inhibition of CKAMP44 primarily downregulates the expression of PPP3r2, which leads to a decrease in its dephosphorylation activity, thereby increasing the phosphorylation level of GluA1 and increasing the membrane expression of GluA1A rescue experiment was conducted in which we effectively inhibited CKAMP44 and overexpressed PPP3r2 simultaneously to validate this conclusion, and we found that overexpressing PPP3r2 reduced the phosphorylation level of GluA1-Ser831 and the membrane expression of GluA1 to exacerbated seizure activity in TLE mice, which antagonized the protective effects of CKAMP44 knockdown.

## Limitations

This study had several limitations. First, we only evaluated the role of CKAMP44 knockdown expression in the KA-induced TLE mice, and future studies should investigate the mechanism of overexpression of CKAMP44 in the KA-induced TLE mice. Second, while CKAMP44 downregulation may affect the expression of multiple downstream proteins, we only examined PPP3r2; thus, further studies should investigate whether and how CKAMP44 is involved in mediating the expression of other proteins.

In conclusion, in KA-induced TLE mice, the inhibition of CKAMP44 attenuated the severity of seizures primarily through the CKAMP44-PPP3r2-GluA1 pathway, thus exerting a protective effect on TLE mice. These findings could provide useful evidence for understanding the role of CKAMP44 in the pathological mechanisms of TLE and developing corresponding clinical therapeutic targets.

## Declarations

### Ethical Approval

Our studies were carried out in strict accordance with the recommendations in the Guide for Chongqing Medical University Experimental Animal Centre. All procedures were approved by the Ethics Committee of Chongqing Medical University, which is fully accredited by the Association for Assessment and Accreditation of Laboratory Animal Care.

### Competing interests

None of the authors has any conflict of interest to disclose. We confirm that we have read the Journal’s position on issues involved in ethical publication and affirm that this report is consistent with those guidelines.

### Authors’ contributions

The research study was conceived and conceptualized by Lihong Huang. Research was designed by Lihong Huang, Shiyu Chen and Haokun Guo. Experiments were performed and analyzed by Lihong Huang, Shiyu Chen, Haokun Guo, Hui Zhang, Xiaoqin Wang, Yiming Guo and Shiyun Yuan. Manuscript draft was written by Lihong Huang and Shiyu Chen, edited by Yang Lü and Weihua Yu. Resources were given by Jing Luo, Liang Wang, Yang Lü and Weihua Yu. All authors have read and approved the final manuscript.

### Funding

This study was supported by grants from the National Natural Science Foundation of China (81871019, 81671286).

### Availability of data and materials

The raw data supporting the conclusions of this article will be made available by the corresponding authors.

## Notes

### Competing Interest Statement

The authors have declared no competing interest.

