## Supplementary material for "Inhibition of CKAMP44 attenuated seizure activity via protein phosphatase 3 regulatory subunit B-mediated GluA1 phosphorylation and synaptic transmission": Appendix A. Supplementary material.docx

**Supplementary material 1**

**Table 1 Reagents of cell culture**

| **Name** | **Catalogue number** | **Company** |
| --- | --- | --- |
| Poly-D-Lysine | A3890401 | Gibco, USA |
| PBS (1X), cell culture grade | MA0015 | meilunbio, China |
| DMEM basic | C11995500BT | Gibco, USA |
| FBS | 10091- 148 | Gibco, USA |
| Penicillin-Streptomycin | 15140- 122 | Gibco, USA |
| Neurobasal (TM) Medium  (1X) | 21103-049 | Gibco, USA |
| B-27 Supplement (50X) | 17504-044 | Gibco, USA |
| Trypsin-EDTA (1X), liquid 0.25% Trypsin with EDTA | 25200-056 | Gibco, USA |
| GlutaMAX Supplement | 35050061 | Gibco, USA |
| D-Hanks | H1045 | Solarbio, China |
| Lipofectamine™ 3000 Transfection Kit | L3000-015 | Invitrogen, USA |

**Table 2 Reagents of protein extraction**

| **Name** | **Catalogue number** | **Company** |
| --- | --- | --- |
| RIPA buffer(high) | R0010 | Solarbio, China |
| Protease Inhibitor Cocktail | HY-K0010 | MedChemExpress, USA |
| Phosphatase inhibitor cocktail D | P1096 | Beyotime Biotechnology,China |
| Enhanced BCA Protein Assay Kit | P0010 | Beyotime Biotechnology,China |
| 5x SDS Loadding Buffer | WB-0091 | Dingguo, Changsheng Biotechnology, China |
| Minute^TM^ Plasma Membrane | SM-005 | Invent, USA |
| Minute™ Denaturated protein solution | WA-009 | Invent, USA |

**Table 3 Reagents of Western blotting**

| **Name** | **Catalogue number** | **Company Dilution** |
| --- | --- | --- |

| Rabbit anti-CKAMP44 | NBP1-76496 | Novus Biologicals, USA | 1:1000 |
| --- | --- | --- | --- |
| Mouse anti-GluA1 | 67717-1-Ig | Proteintech, China | 1:1000 |
| Rabbit anti-GluA2 | 11994-1-AP | Proteintech, China | 1:1000 |
| Rabbit anti-GluA3 | #4676 | Cell Signaling Technology, UK | 1:1000 |
| Rabbit anti-GluA4 | #8070 | Cell Signaling Technology, UK | 1:1000 |
| GluA1 phospho S831 | ab109464 | Abcam | 1:1000 |
| GluA1 phospho S845 | 381354 | zenbio | 1:1000 |
| Rabbit anti-PPP3r2 | 14005-1-AP | Proteintech, China | 1:1000 |
| Rabbit anti-GAPDH | 10068- 1-AP | Proteintech, China | 1:5000 |
| Rabbit anti-calnexin | 66903-1-lg | Proteintech, China | 1:2000 |

| HRP Goat Anti-Rabbit IgG(H+L) | SA00001-2 | Proteintech, China | 1:5000 |
| --- | --- | --- | --- |
| Protein free fast blocking solution (5 x) | PS108 | Beyotime |  |
| Tris | 77-86- 1 | GENVIEW, USA |  |
| Glycine | 56-40-6 | GENVIEW, USA |  |
| SDS | GS286 | GENVIEW, USA |  |
| NaCl | 7647- 14-5 | GENVIEW, USA |  |
| TWEEN® 20 | 9005-64-5 | Sigma-Aldrich, USA |  |
| PAGE Gel Fast Preparation Kit | PG111 | Epizyme Biomedical |  |
| (7.5/10/12.5%) |  | Technology, China |  |
| Ultrasensitive ECL Western HRP  Substrate [Low-fg Level] | 17047 | ZENBIO, China |  |

| **Table 3 Reagents of real-time quantitative PCR** | | |
| --- | --- | --- |
| **Name** | **Catalogue number** | **Company** |
| SimplyP Total RNA  Extraction Kit | BSC52S1 | BioFlux, China |
| HiScript® II Q RT SuperMix for qPCR (+gDNA wiper) | R223-01 | Vazyme, China |
| ChamQ Universal SYBR qPCR Master Mix | Q711-02 | Vazyme, China |
| RNase-free ddH2O | P071-01 | Vazyme, China |

**Table 4 Reagents of immunofluorescence staining**

| **Name** | **Catalogue number** | **Company** | **Dilution** |
| --- | --- | --- | --- |
| Rabbit anti-CKAMP44 | NBP1-76496 | Novus Biologicals, USA | 1:1000 |
| Mouse anti-NeuN | 66836-1-lg | Proteintech, China | 1:200 |
| Mouse anti-GFAP | 60190-1-lg | Proteintech, China | 1:300 |
| CoraLite488  Anti-Rabbit IgG (H+L) | SA00013-2 | Proteintech, China | 1:500 |
| CoraLite594  Anti-Mouse IgG (H+L) | SA00013-3 | Proteintech, China | 1:500 |
| DAPI | 0100-20 | SouthernBiotech,  USA |  |

**Table 5 Virus construction**

| **Name** | **Titer** | **vector** | **Company** |
| --- | --- | --- | --- |
| AAV-shisa9 | 1. 17E+09 TU/mL | GV478-U6-MCS-CAG-EGFP | Genechem Corp.,Ltd.  (Shanghai, China). |
| AAV-Control | 2.61E+08 TU/mL | CON305-U6-MCS-CAG-EGFP | Genechem Corp.,Ltd. (Shanghai, China). |
| AAV-PPP3r2 | 1.3* 10^ 12 vg/mL | pAAV-CMV-PPP3r2-3xFLAG-  P2A-mCherry-WPRE | Obio Technology Co., Ltd. (Shanghai, China). |
| AAV-Control | 1.0* 10^ 12 vg/mL | pAAV-CMV-MCS-3xFLAG-P2A-mCherry-WPRE | Obio Technology Co., Ltd. (Shanghai, China). |

TU：Transducing Units

vg：Vector Genomes

**Table 6 RNA primer of qPCR**

| **Gene**  **name** | **Forward primer** | **Reverse primer** |
| --- | --- | --- |
| Mouse | 5'-CAGTGGCAAAGTGGAGATTGTTG- | 5'-TCGCTCCTGGAAGATGGTGAT-3' |
| *Gapdh* | 3' |  |
| Mouse | 5'-CACCTATGAGCCCGACCCTATTA- | 5'-TTTCACCAACAGTCTCAGCTTGA- |
| Shisa9 | 3' | 3' |
| Mouse | 5'-AACTGAGCTGTGCAACCACT- | 5'-TTGCCGTCTGTGTCGAAGAT- |
| *Ppp3r2* | 3' | 3' |
