## Supplementary figures and images for "Inhibition of CKAMP44 attenuated seizure activity via protein phosphatase 3 regulatory subunit B-mediated GluA1 phosphorylation and synaptic transmission"

### Appendix Figure 1.png

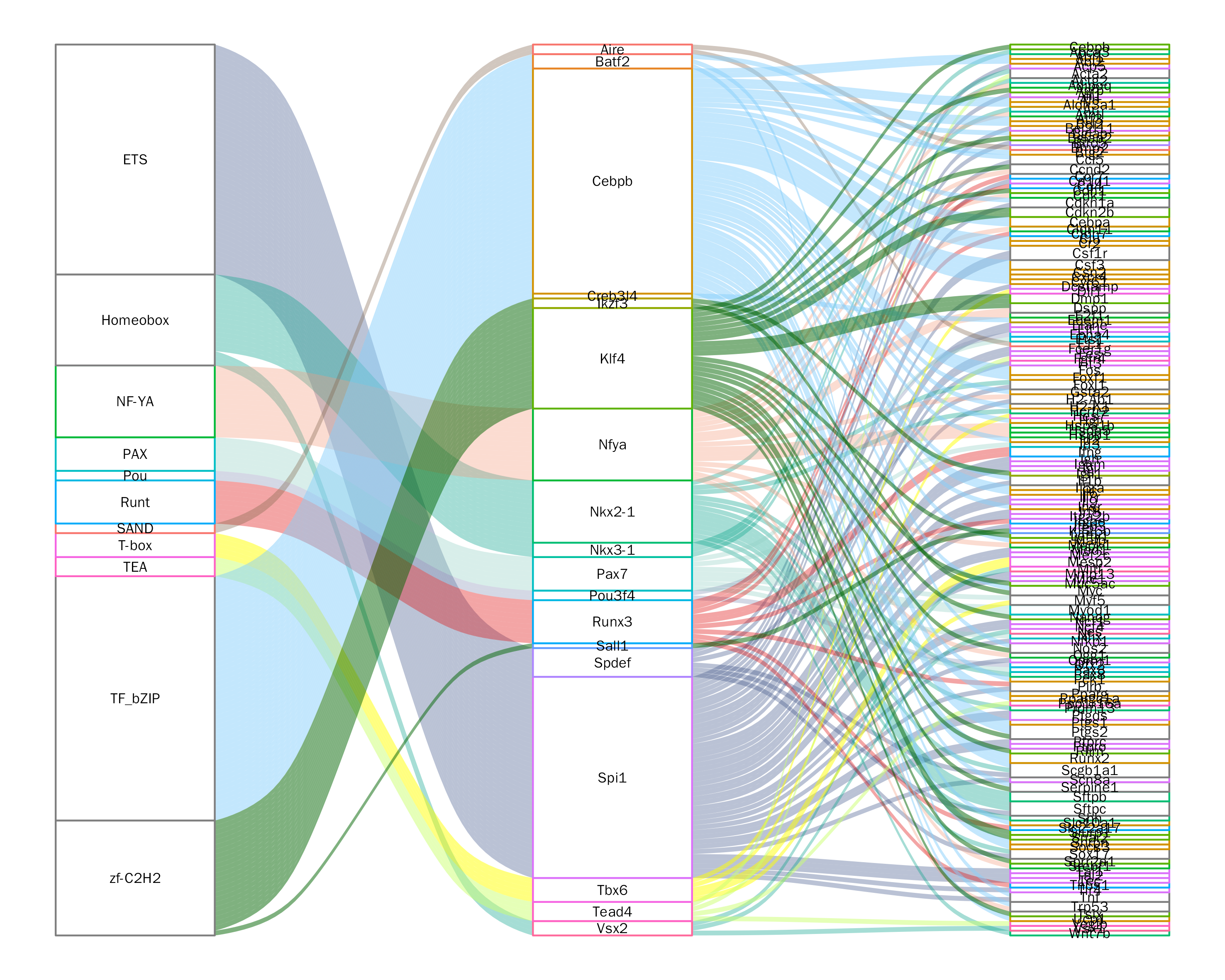
